# Competition between memories for reactivation as a mechanism for long-delay credit assignment

**DOI:** 10.1101/2025.11.08.687292

**Authors:** Subhadra Mokashe, Paul Miller

**Author notes:** **For correspondence:** (PM).

## Abstract

Animals can associate events with their outcomes, even if there is a long delay between the two. For example, in conditioned taste aversion, animals gain an aversion to a taste (the conditioned stimulus, CS) if sickness (an unconditioned stimulus, US) is induced up to 12 hours later. Established correlational plasticity mechanisms, operating on timescales of milliseconds to seconds, do not wholly explain how networks of neurons achieve such long-delay credit assignment. Moreover, if the animal experiences an intervening taste (an interfering stimulus, IS), the IS gains some “credit” for the causality of the outcome, reducing aversion to the CS. We hypothesize that reactivation of prior events at the time of outcome causes specific associative learning between those events and the outcome. We explore the inherent competition underlying credit assignment using a spiking neural network model storing memories through time-decaying synaptic strengthenings in two groups of neurons producing inherently competing attractor states. We show how the later memory can be reactivated more often and reduce the reactivation of a prior memory. Also, we provide a mechanism for the experimental finding of a rebound in association with, and therefore aversion to, the CS if the time between the following IS and US is increased. Such a result can appear paradoxical as associations typically diminish with time, but arises when the IS initially produces a strong decrease in reactivation of the CS, but reactivations of the CS thereafter increase, in spite of weakening synaptic strengths, because competing reactivations of the IS decrease more. By reactivating the memories probabilistically, neural circuits can assign the credit in a biologically plausible way.

## Introduction

Animals have evolved the remarkable ability to learn from new experiences and to associate the value of any outcome of that experience with the actions preceding it. This allows us to draw from past experiences to optimize our actions by maximizing those positive outcomes that enhance our likelihood of survival. Such associative learning is orchestrated by synaptic plasticity during learning, which changes connections in neural circuits such that when we experience the same action or stimulus again, we can also recall the outcome. Synaptic plasticity in the connections between neurons is engaged when they are active close in time. Therefore, synaptic plasticity can account for the association of an action with its outcome when the outcome immediately follows the action, as neuronal assemblies separately representing action and outcome would be active close in time, and the connections from neurons representing action to neurons representing outcome could be strengthened. However, the mechanisms underlying the temporal credit assignment become elusive when the action and the outcome are separated by a significant delay (***Izhikevich, 2007; Päpper et al., 2011***). One such long-delay association task is conditioned taste aversion (CTA), in which an animal, when exposed to a novel taste in the hours preceding induced sickness, develops an aversion to that taste (***Nachman, 1970; Adaikkan and Rosenblum, 2015***).

Reactivation of neural activity in cells that responded to prior stimuli is seen in many regions, including the hippocampus (***Ólafsdóttir et al., 2018; Diekelmann et al., 2012; Carr et al., 2011***), the sensory cortices (***Ji and Wilson, 2007; Barnes and Wilson, 2014***), as well as the amygdala (***Sara, 2000***). Such reactivation is likely relevant in the gustatory cortex (GC) and/or the basolateral amyg-dala (BLA), as CTA depends on a coordination between these two brain regions (***Naor and Dudai, 1996; Grossman et al., 2008***). Reactivation of the neural representation of the prior conditioned stimulus (CS) at the time of sickness could lead to the neurons representing taste being active at the same time as neurons indicating sickness. Such simultaneous activity would enable synaptic plasticity to strengthen connections in a manner that forms an association between the CS and sickness. Indeed, recent evidence suggests that reactivation of neurons in amygdalar networks during malaise helps associate a novel flavor with sickness (***Zimmerman et al. (2025)***).

During the long time delay between an action and its outcome, an animal could perform other actions or sample other stimuli, making temporal “credit assignment”-the term used for associating action and corresponding outcome, even if the latter is negative-a difficult problem. For example, in taste aversion, if a second substance is tasted between the first and the malaise, it is not completely clear which taste caused the malaise. The difficulty is solved in part because aversion is toward novel tastes–a substance that has been consumed often in the past would not be considered harmful after one bad experience. In the case of two novel tastes, when the second tastant —an interfering stimulus (IS) —is sampled between the CS and sickness induction, aversion to the CS is reduced compared to when only the CS is sampled by the animal. Moreover, the animal gains an aversion towards the IS, which is greater than that it now has to the CS. The reduction in association of the CS with sickness, due to the introduction of an IS, is known as overshadowing (***Schachtman et al., 1992; Revusky, 1971***). We, like many others, assume that overshadowing is the outcome of a competition between stimuli for association with the sickness. We hypothesize that the neural representations of the two stimuli compete for reactivation during sickness, and the competition shapes the extent to which the IS overshadows the CS as the cause of sickness.

A recent study ***Kwok et al. (2017***) characterized how the timing of the IS affects the degree of aversion towards both the CS and the IS. A later IS shows more overshadowing compared to an earlier IS when the interval between CS and sickness is held constant. Presumably, this is a reflection that the later IS, being closer in time to the sickness, appears to be more significant and more likely to have caused the sickness. Such a result is in general alignment with the finding that the strength of aversion to a sampled taste gradually diminishes over many hours as the time between taste and sickness increases (***Adaikkan and Rosenblum, 2015***). If overshadowing is considered as a competition, then a later IS means less time between IS and malaise, so a stronger aversion to the IS and therefore a weaker aversion to the CS. The next set of experiments produced an additional intriguing result. When the time between CS and IS was held fixed, but the delay to sickness onset was increased, the aversion to the CS actually increased, showing a greater association between stimulus and outcome in spite of a greater delay between the two. Effectively, the overshadowing had decreased, as the delay between IS and sickness had become more similar (proportionally) to the delay between CS and sickness. One could think of the ratio of times between stimuli as outcome being an indicator of which is more likely, such that the longer delay actually makes the earlier stimulus more likely to be a cause than when the IS is comparatively much closer to the sickness. In this study, we aim to understand if competition for reactivation between the neural representations of the two stimuli could explain these results.

Our model shows how the competition between assemblies for reactivation could explain the overshadowing of the CS by the IS. We also study the effect of the inter-stimulus interval, and our results could account for the finding that a later IS shows more overshadowing when the CS to sickness interval is held fixed. On changing the delay to the outcome, we find that for a specific decay profile of the recurrent strengths, the animals could experience a rebound in conditioning towards the CS, as is seen ***Kwok et al. (2017***), for the reasons stated above. Therefore, we can provide an account for overshadowing and its dependence on the relative timing of two stimuli and an outcome by studying the competition for reactivation of neural assemblies–which could be thought of as competing memories– and establishing a mechanism for resolving temporal credit assignment ambiguities.

## Results

### Reactivation of memories

We start by testing our hypothesis that competition for reactivation shapes credit assignment by building a spiking network model with clustered connectivity (Figure 1B). The strength of aversion decreases as the delay between the taste and the sickness increases (***Adaikkan and Rosenblum, 2015***). To account for that, we assume the recurrent weight of the CS and the IS ensembles decreases with time (Figure 1 A). Their values at the time of sickness induction are *a* (CS) and *a* + Δ (IS), respectively. IS has a higher recurrent weight as it was introduced later. The time between the IS and sickness is less, so there is less decay (Figure 1 A, B). All neurons receive independent Poisson inputs, but all the excitatory neurons share the rate of the Poisson processes, which is given by an OU process (See Figure 1 C). We employ an OU process for rate modulation, allowing temporally extended input upticks that trigger reactivation events in the network (Figure 1D).

**Figure 1.**
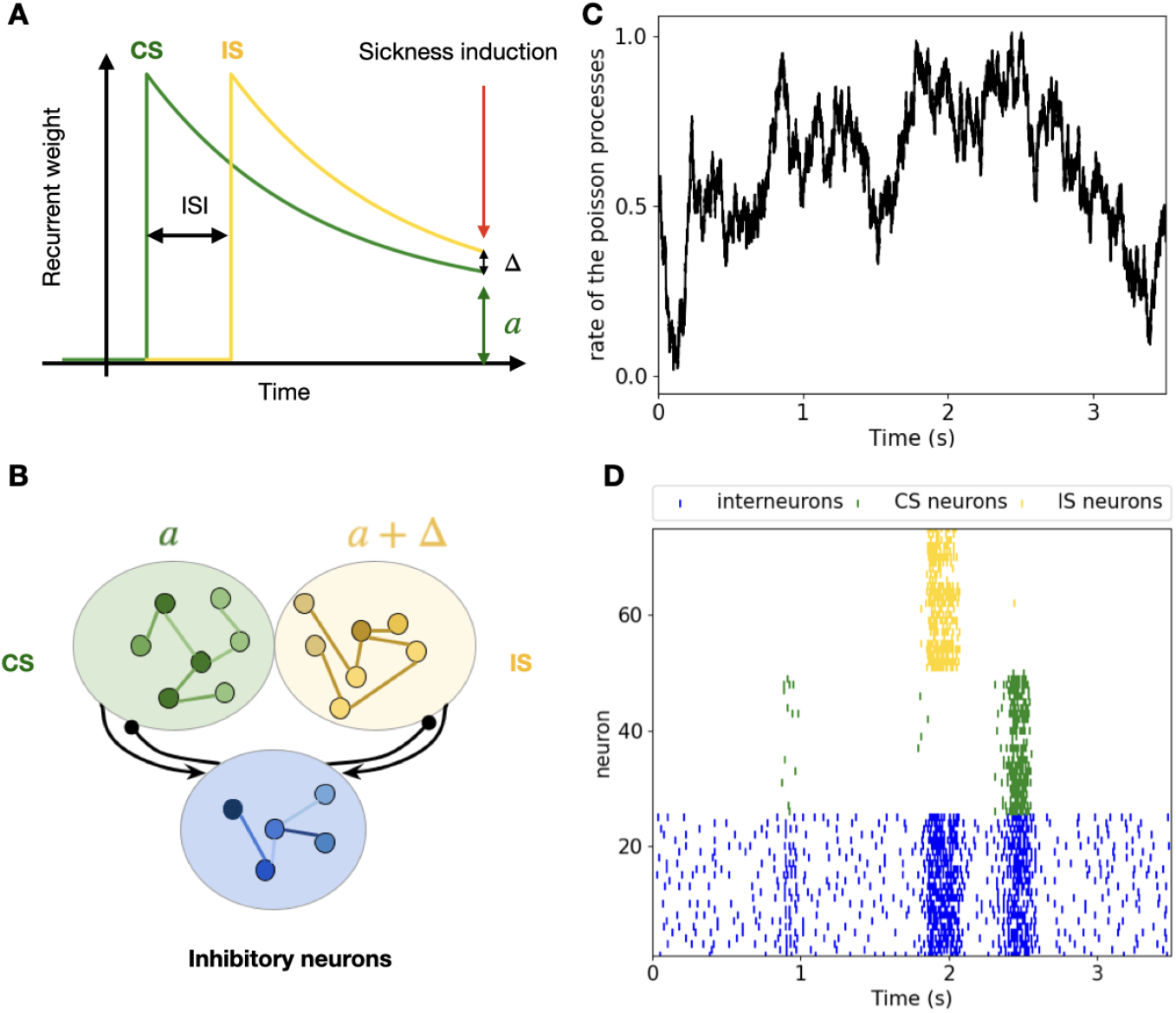
A. The assumed decay of recurrent weights gained after activation of the taste representation. Note the higher recurrent strength of the IS ensemble in A and B. B. Illustration of the network structure. C. The OU process shared by all excitatory neurons shapes the rate of the independent Poisson processes. D. Raster plot of a fraction of neurons from each pool showing reactivations. Note that either the CS or the IS ensemble is active at a given time.

### Preferential reactivation of a more recent input

First, we want to see how changing the inter-stimulus interval affects the aversion to the tastes (Figure 2A). We assume that the more time the network spends reactivating a taste, the greater the aversion to that taste (Figure 2B; notice the almost equal amount of time spent reactivating the CS and the IS at Δ = 0). We see that on increasing the inter-stimulus interval or the Δ, the network spends more time reactivating the IS, and we see a reduction in the time spent in the CS state (Figure 2 C). This implies that a later IS would have more conditioning towards the malaise, which is exactly what is seen in the overshadowing experiments in ***Kwok et al. (2017***). Thus, our model can capture that a later IS would have stronger conditioning towards the US and would show more overshadowing. We also observe a decrease in time spent in the CS state, which would lead to a lower association with malaise and matches the lower conditioning seen towards the CS, explaining the overshadowing of the CS by the IS (Figure 2 C).

**Figure 2.**
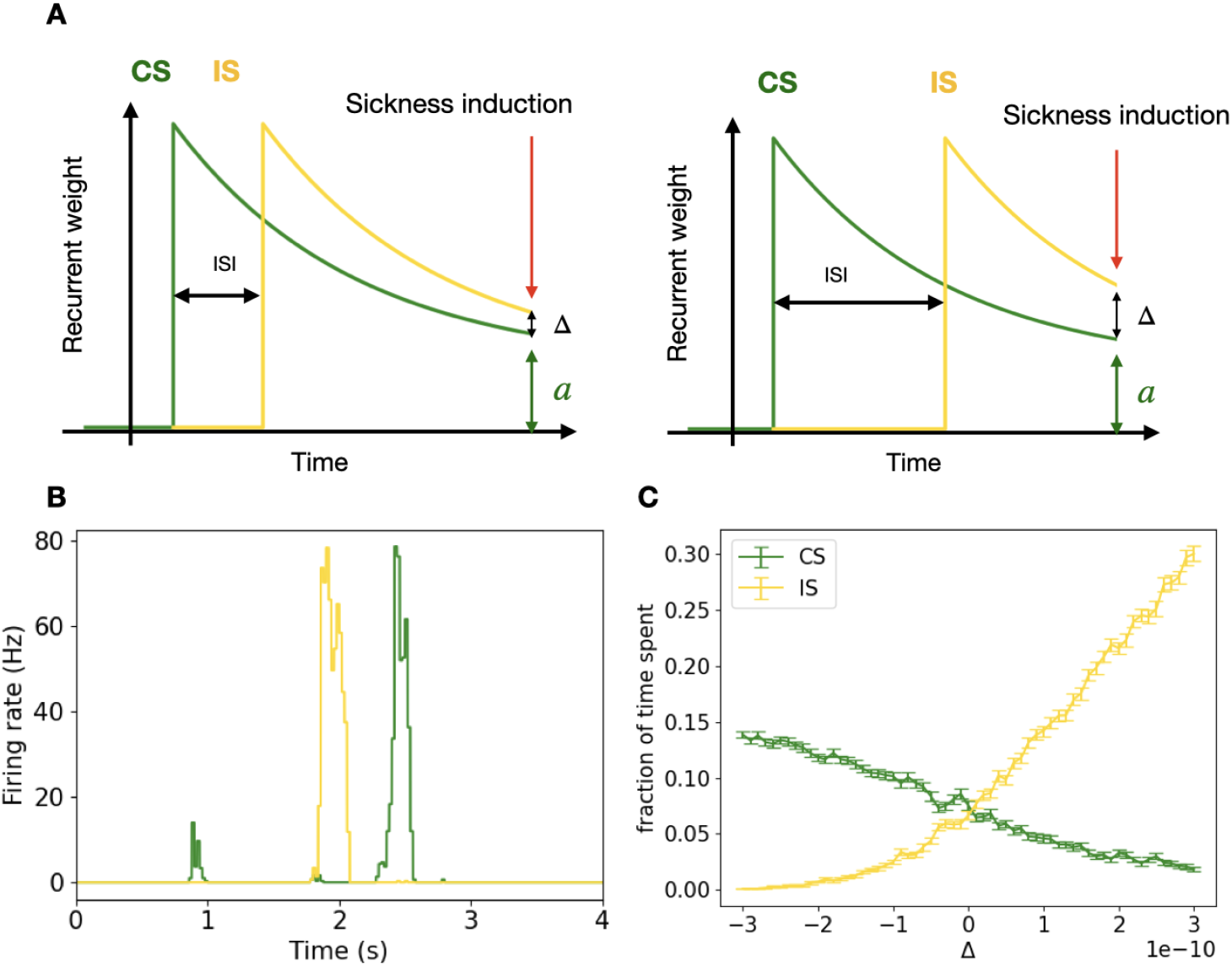
Reactivation as a function of Δ: A. Illustration of how changing the inter-stimulus interval changes the value of Δ. B. Reactivations in network simulations at Δ = 0. C. The fraction of time the network spends reactivating IS and CS states. Note the decrease in the time spent in CS as a function of Δ.

The time spent in the IS increases as a function of Δ as the recurrent strength of the IS ensemble increases. On the contrary, on increasing the Δ, the recurrent strength of the CS ensembles remains unchanged, yet we see a reduction in the time spent by the network in the CS (Figure 2 C). If the fraction of total time spent in either CS or IS was close to one, such a reduction in the time spent in CS could be chalked up to the lack of time for the reactivation of the CS. When the fraction of time spent in either CS or IS is around 0.15, we see that the correlated noise, rather than competing for time, drives the reduction in the time spent in the CS state. The competition is at its lowest when the ensemble rates are determined by two independent OU processes (competition index = -0.0422; see Methods for the exact definition of the competition index). When we have 25% noise correlation, we see an increase in competition (competition index is -0.05), and at 50% correlation, the competition index is -0.07.

When the rate of fluctuations is completely correlated, we observe high competition (the competition index is -0.1658). The noisy inputs received by the CS and the IS ensemble follow the same OU process as the Poisson input rate. Having the same OU process dictate the rate of the Poisson process for all excitatory neurons (both ensembles) results in limited upticks in fluctuations when reactivations could occur, making the firing sparse, as seen in the gustatory cortex (Unpublished data by Hannah Germaine, Katz lab). Limiting the time during which the neurons fire effectively turns it into a time-based competition. As correlation increases, the likelihood of both ensembles being activated together increases. Both ensembles activate the inhibitory neurons. The ensemble with the higher recurrent weight receives more recurrent inputs and is activated more often. This increases inhibition of the other ensemble, suppressing reactivation. When there is little or no correlation between the inputs, the likelihood that both ensembles are activated together decreases, reducing competition between the inputs.

### Late sickness onset causes less overshadowing

***Kwok et al. (2017***) show that late sickness leads to less overshadowing than early sickness, i.e., greater conditioning towards the CS for late sickness than for early sickness. Such a rebound in conditioning towards the CS is puzzling, as without the presence of an IS, conditioning towards the CS decays over time (***Adaikkan and Rosenblum, 2015***), as shown in Figure 3 D (black line). We propose that a specific decay profile of recurrent strengths in the network can account for the observed rebound in CS conditioning. A sum of two exponential functions where one time constant is much longer than the other leads to a rapid decay of the difference between the two traces compared to the recurrent strength (Figure 3 A). We calculate the time spent in the CS state for each value of recurrent strength and delta. As expected, time spent in CS decreases as a function of Δ and increases as a function of *a* (Figure 3 B, C). We then look at the time spent in CS along the *a*&Δ trajectory from Figure 3 A and find a rebound in the time spent in the CS state by the network (Figure 3 D). This rebound is seen even though the recurrent strength for CS is decaying in time (Figure 3 A). This rebound in time spent in CS would explain why later sickness induction causes less overshadowing. To get the rebound, a specific set of decay functions for the recurrent weight is needed, and we get the paradoxical result that even though the recurrent strengths within the CS ensemble are weakened, that ensemble reactivates more frequently because the recurrent strengths within the IS ensemble are weakened even more with greater decay and produce less competition.

**Figure 3.**
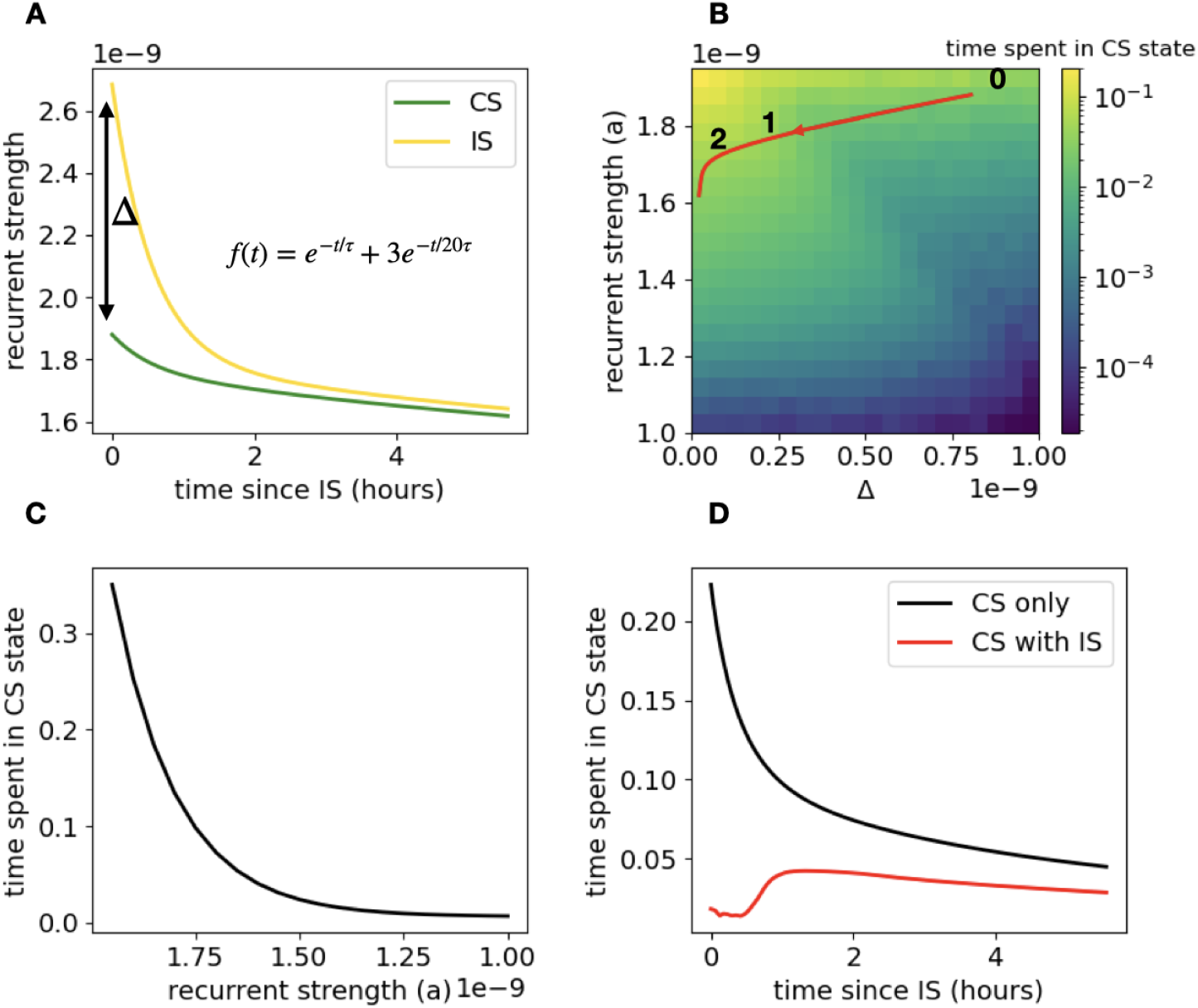
Rebound in conditioning. A. Recurrent weights decay with time. Note the difference between the two traces given by Δ reduces much faster than the green trace itself. B. The same trajectory as A in the *a*, Δ space. C. Trajectory taken by the recurrent weight in units of recurrent strength *a* when no IS is present. D. Time spent in CS along the trajectory when IS is presented and when no IS is presented. Notice the rebound in the time spent in the CS at ∼ 1 hour.

### Competition as a function of recurrent strength

We change the recurrent strength of the CS to simulate a change in the delay to the sickness onset. As the onset of sickness is delayed, the recurrent strengths of both the IS and CS ensembles decrease, as does the difference between their recurrent strengths (Figure 4 A). A non-monotonic relationship between the competition index and recurrent strength *a* arises out of the existence of bistability only at intermediate values of *a* as a function of the current amplitude (Figure 4 B) . At higher values of *a*, both ensembles can be active, reducing competition. At lower values of *a*, neither ensemble is sufficiently activated to inhibit the other.

**Figure 4.**
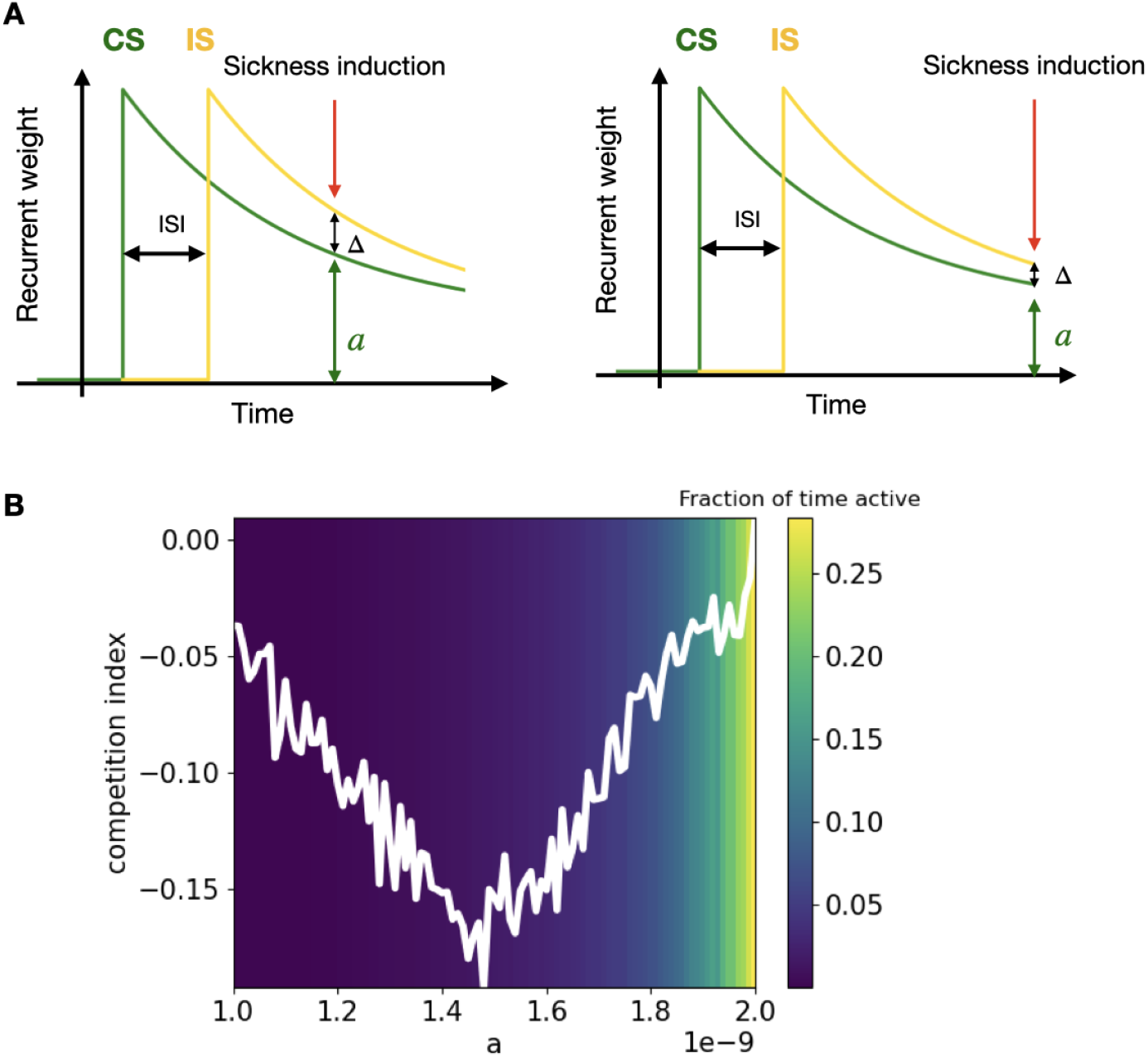
Reactivation as a function of *a*: An illustration of how changing the delay to sickness onset changes the value of *a*. B. Competition index as a function of *a*.

**Figure 5.**
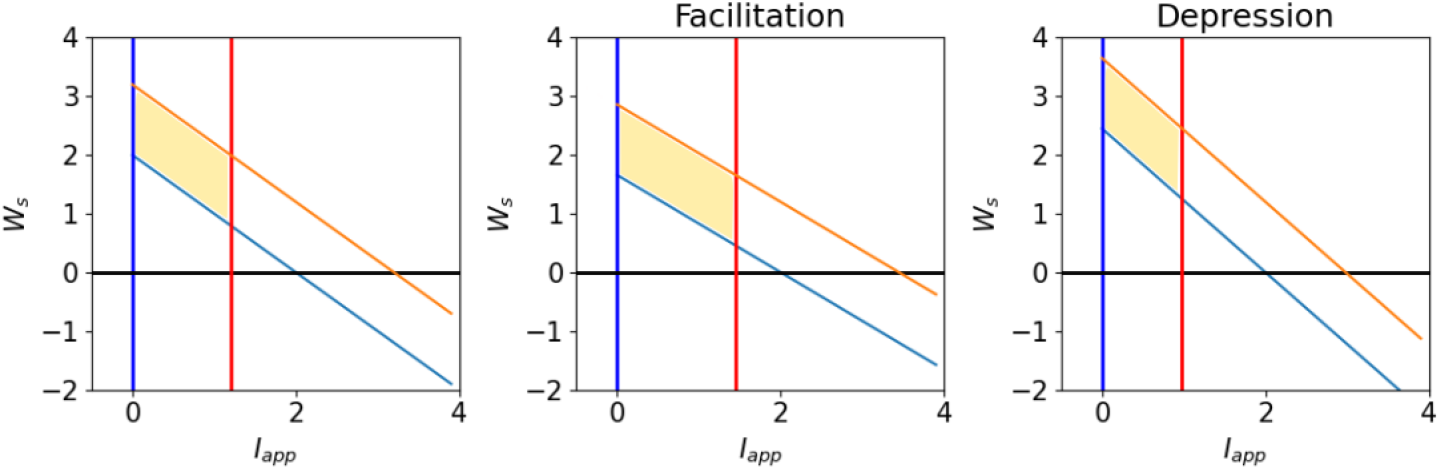
Bistability regions (shaded area) as a function of the recurrent weight and current amplitude for facilitation and depression. Notice the expansion of area in the case of facilitation and contraction of area in the case of depression, compared to the case with no short-term plasticity.

### Bistability

Facilitation of excitatory synapses causes the recurrent strengths of the ensembles to transiently increase, mimicking a change in the recurrent strength (*a*). As correlated noise causes the competition, facilitation in the E to E connections would increase the competition, as the ensemble that is being reactivated will get further strengthened, blocking the other ensemble from being activated. Depression of synapses would have the opposite effect. We model short-term synaptic plasticity to study its effect on the degree of competition between the two memories. We want to understand how depression and facilitation of synapses change the degree of competition between the ensembles.

For competition to occur, we need the network to be in a bistable regime—where the active states of the CS or the IS are stable. [*CS, I S*] = [1, 0] or [0, 1], where 1 is active and 0 is inactive, should be the stable states, called reactivation states. Fluctuations that cause a reactivation event activate both ensembles; then, inhibition kicks in, and the ensemble with the higher net current wins. This occurs in the bistability region as a function of *a* and *I*_*ext*_. As the current amplitudes fluctuate with noise, the larger the bistability region, the more likely it is that current fluctuations will lie within it. The larger the bistability region, the greater the competition.

To study how bistability changes with facilitating and depressing synapses, we study a reduced model of the network where the rates of the two units *r*_1_ (CS) and *r*_2_ (IS) are given by the following equations:

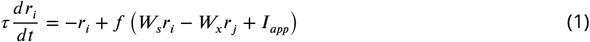

We assume that CS and IS are presented simultaneously. *W*_*s*_ is the recurrent strength and *W*_*x*_ is the cross inhibitory weight, *I*_*app*_ is the applied current and *f* (*I* ) is the input-output function.

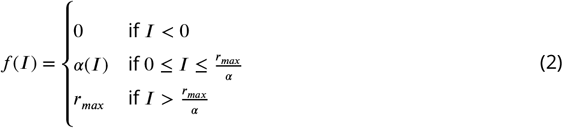

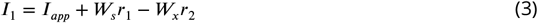

We are interested in finding the regions of bistability in terms of the parameters *W*_*s*_ and *I*_*app*_. We derive a closed-form expression for the area of the bistable region. We also find the areas for the region of bistability in the case of depression and facilitation. The larger the area where the network is bistable, the more likely it is that a current fluctuation will put the network into the bistable regime. The larger the bistability region, the greater the competition. To find the regions of bistability, we want to identify regions where unit 1 is stable at *r*_*max*_ and unit 2 is stable at 0, or where unit 1 is stable at 0 and unit 2 is stable at *r*_*max*_. Since the two units are symmetric, we can focus on the conditions on *I*_1_. For analytical tractability, we assume the active unit’s rate is *r*_*max*_ while the inactive unit’s rate is zero. We will now derive the area of the bistability region.

For the point [*r*_1_, *r*_2_] = [0, 0] to be unstable, i.e. the condition for *r*_1_ ≠ 0. For *r*_1_ ≠ 0, given the transfer function we need,

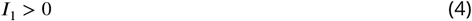

at [*r*_1_, *r*_2_] = [0, 0], *I*_1_ = *W*_*s*_ × 0 + *W*_*x*_ × 0 + *I*_*app*_.

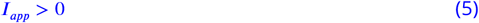

For the stable point at [*r*_1_, *r*_2_] = [*r*_*max*_, 0], i.e. the condition for *r*_1_ = *r*_*max*_

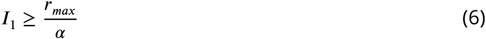

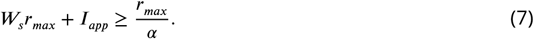

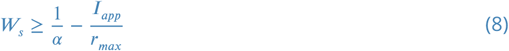

For the stable point at [*r*_1_, *r*_2_] = [0, *r*_*max*_], the condition for *r*_1_ = 0

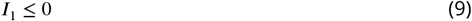

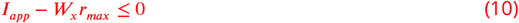

For the unstable point at [*r*_1_, *r*_2_] = [*r*_*max*_, *r*_*max*_]

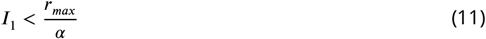

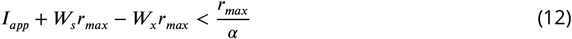

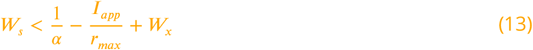

As the region enclosed by the lines is a parallelogram, we can find it’s area: *W* ^2^*r*_*max*_.

In the case of facilitation,

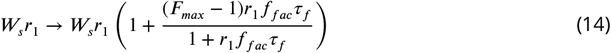

For the unstable point at [*r*_1_, *r*_2_] = [0, 0], i.e. condition for *r*_1_ ≠ 0

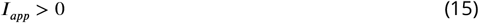

For the stable point at [*r*_1_, *r*_2_] = [*r*_*max*_, 0], i.e. condition for *r*_1_ = *r*_*max*_

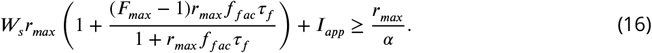

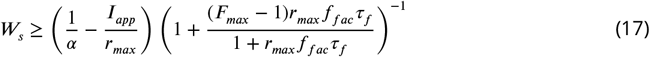

For the stable point at [*r*_1_, *r*_2_] = [0, *r*_*max*_], the condition for *r*_1_ = 0

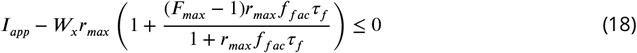

For the unstable point at [*r*_1_, *r*_2_] = [*r*_*max*_, *r*_*max*_]

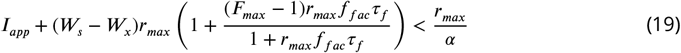

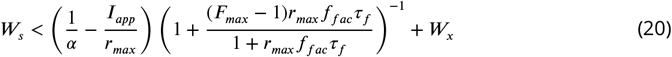

Area with bistability 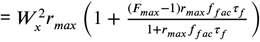

As 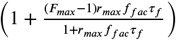 is greater than one, the area of the region of bistability increases in the case of facilitation.

In the case of depression,

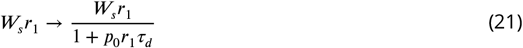

For the unstable point at [*r*_1_, *r*_2_] = [0, 0], i.e. condition for *r*_1_ ≠ 0

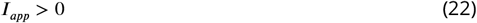

For the stable point at [*r*_1_, *r*_2_] = [*r*_*max*_, 0], i.e. condition for *r*_1_ = *r*_*max*_

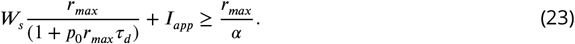

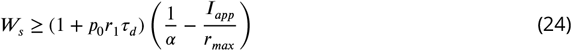

For the stable point at [*r*_1_, *r*_2_] = [0, *r*_*max*_], the condition for *r*_1_ = 0

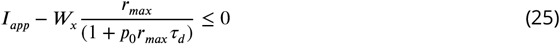

For the unstable point at [*r*_1_, *r*_2_] = [*r*_*max*_, *r*_*max*_]

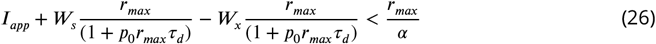

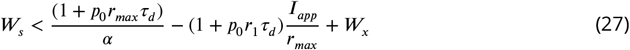

Area with bistability 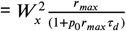

The area of the parallelogram increases with facilitation as area 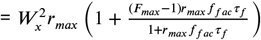 which is greater than 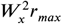 as 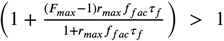 for positive values of 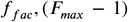 Similarly, for depression the area of the parallelogram decreases as area 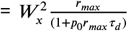 where compared to the area of 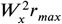. The area of bistability increases with facilitation and decreases with depression, suggesting higher competition with facilitating synapses and lower competition with depressing synapses. Our network simulations align with the theoretical results.

In summary, we have shown that the competition for reactivation could explain the phenomenon of overshadowing. We outline how the credit can be assigned through the competition for reactivation when there is ambiguity about the cause of the outcome.

## Discussion

A growing body of experimental studies of long-delay aversive learning (***Zimmerman et al. (2025); Zaki et al. (2025); Puhger et al. (2024***)) suggests that reactivation of the neurons that respond to the stimulus could help associate a stimulus with a temporally distal aversive outcome. Reactivation, or replay, is seen after learning and has been known to strengthen traces of the resulting memories (***Ólafsdóttir et al., 2018***). Reactivation is also theorized to help integrate new information into existing knowledge structure through consolidation (***Rasch, 2018***) and in the hippocampus, is proposed to associate discontiguous events (***Wallenstein et al., 1998***). We hypothesize that reactivation helps associate temporally separated events, particularly when there are long delays between the events.

How credit is assigned when more than one event could be the cause of an outcome is an active area of research (***Witkowski et al., 2025; Izhikevich, 2007***). We investigate the temporal credit assignment in the case of overshadowing in taste aversion learning when two temporally separated stimuli are sampled by the animal, before it is made sick by a LiCl injection (***Schachtman et al., 1992***). The animal shows higher aversion towards the interfering stimulus (IS), which is sampled later, and shows less aversion towards the conditioned stimulus (CS) than it does in experiments with the CS and no IS. We propose that reactivation helps associate the stimuli with the sickness, and competition for reactivation between assemblies of neurons representing the two stimuli could shape the relative degree of aversion seen towards each stimulus. If such differences in learned aversion are considered differences in memory, then our model is one of competition between memories.

We show that competition between neural assemblies for reactivation arises out of the limited number of opportunities when the fluctuating excitatory input to the circuit is large enough to trigger a reactivation event. Such competition depends on the two neural assemblies, representing the CS and the IS, receiving correlated fluctuations. During these upticks, both assemblies are excited and begin to activate, which in turn further activates the inhibitory pool of neurons. While inhibitory neurons inhibit both the CS and IS ensembles, the ensemble with the higher internal connection strengths is more likely to reactivate strongly and suppress the other ensemble. Therefore, over time and multiple reactivation events, the more strongly interconnected ensemble is reactivated more often and therefore able to undergo more associative plasticity with neurons representing the outcome. We also study how inherent dynamical features of synapses – short-term facilitation and short-term depression– affect the competition between the two ensembles and find that facilitation increases while depression decreases competition.

An assumption in our model is that at the time of stimulus experience, the coactive neurons retain greater excitability–for example, due to correlational plasticity at the time of their activation– and that excitability decays over time. In this scenario, since the IS is introduced after the CS, neurons responsive to the IS are more excitable at the time of the sickness, as their enhanced excitability has had less time to decay. We implement such enhanced excitability as an increase in internal connection strength, but other mechanisms (enhanced intrinsic excitability or stronger afferent connections) are also possible. The more excitable assembly of neurons at the time of sickness is reactivated more often and therefore can form a stronger associative memory. Moreover, reactivation of the IS can lead to overshadowing of the CS by the IS because of the ensuing competition described in the previous paragraph. The idea of competition between stimuli has been explored in aversive conditioning (***Herrera et al., 2022; Thorwart et al., 2012; Cannon et al., 1985***) but not through the lens of competition for reactivation. Here, we explore the factors that influence the competition between neural assemblies for reactivation–which can be considered a competition between memories–in long-delay association.

We first examine how the interstimulus interval (the interval between the CS and the IS) changes the degree of overshadowing seen towards the CS, and in doing so, we account for the results seen in experimental studies (Figure 2). It is seen that a late IS shows more overshadowing than an earlier one (***Kwok et al., 2017; Cannon et al., 1985; Kaye et al., 1988***) when the time interval between the CS and outcome is held constant. In the experimental study ***Kwok et al. (2017***), if the IS is introduced 50 min after the CS, it causes more overshadowing (i.e., there is less aversion towards the CS) than when the IS is introduced 10 min after the CS. The results of our simulations account for such a finding, as we see less reactivation of the neural assembly representing the CS when neurons in the assembly representing the IS have stronger internal connections (higher Δ) due to a later IS, compared with trials when those connections are weaker, representing an earlier IS. In experiments where the CS and the IS are presented at the same time, changing the salience of the IS has a consequence equivalent to changing the time of the IS in other experiments ***Lindsey and Best*** (***1973***). When the salience of the two stimuli is comparable, there is reciprocal overshadowing seen between the two stimuli (***Bond, 1983***). Our model would account for such behavior if we assume the more salient the stimulus, the greater its lingering excitability after stimulus offset.

We treat the neural representations of sensory inputs as non-overlapping for simplicity of modeling and analytical tractability in this work. However, such a simplification is generally far from the truth as neural representations of stimuli, for example, those of different taste stimuli in gustatory cortex (***Fletcher et al. (2017); Katz et al. (2002); Mukherjee et al. (2019); Fontanini and Katz*** (***2006***), unpublished data from Hannah Germaine, Katz lab) are generally distributed and overlapping. Interference from correlated memories is one of the primary contributors to forgetting (***Chanales et al., 2019; Sissons et al., 2009***). We study how the overlap between the neural assemblies representing the CS and the IS impacts the degree of competition. We expected low levels of overlap to increase the competition for reactivation of the two assemblies if synapses were depressing and spike rate adaptation was strong within neurons. In such a situation, the shared neurons, after being active in one ensemble, would be less excitable and produce less input, thus reducing the activity of the other ensemble for a short time afterwards. However, we only see a slight increase in competition for very low levels of overlap (when one neuron out of 500 is shared across the two ensembles), but for most values of overlap, the competition is lower than in the case of no overlap. Moreover, with high overlap (i.e., a high percentage of neurons shared across assemblies), the two assemblies reactivate together. Inter-stimulus associations are indeed shown to reduce overshadowing (***Sissons et al., 2009***) and agree with our finding of less overshadowing with greater overlap between assemblies.

Our simulations account for a second intriguing experimental finding ***Kwok et al. (2017***). If the time between CS and IS is held constant (at 50 mins) and the time to sickness onset is increased (from 65 mins after the CS and 5 mins after the IS to 95 mins after the CS and 35 mins after the IS) then while aversion to the IS decreases as expected with the greater delay, the aversion to the CS surprisingly increases. Accounting for such a rebound —by observing a greater association between CS and malaise with increased delay between the two —was one goal of our study. We find that for a specific temporal decay profile of recurrent weights, initial fast decay followed by slower decay, as could arise from different biochemical mechanisms underlying synaptic plasticity (***Adaikkan and Rosenblum, 2015; Igaz et al., 2004***), we do indeed observe a rebound in the time spent in the CS (Figure 3). Our result relies on the strong competition between CS and IS, and that while the excitability of the ensemble representing the CS is always decaying, the initially faster decay in excitability of the assembly representing the IS and resulting reduction in its competition for reactivation can have a larger impact on the assembly representing the CS, allowing it to reactivate more often in spite of its reduced excitability.

Although we focus on overshadowing in conditioned taste aversion, overshadowing occurs in conditioning experiments dependent on many different sensory modalities and can occur across sensory modalities (***Pinto and Pineida, 2020; Protzko et al., 2023***). A more salient (strong) odor overshadows a less salient odor when a mixture of odors is presented during olfactory aversion paradigms (***Schubert et al., 2015; Tovar-Díaz et al., 2011***). Overshadowing is seen in the interaction of visual and verbal inputs, where the verbal description of visual inputs impairs one’s ability to later recognize or recall that visual information (***Schooler and Engstler-Schooler, 1990***). Serial over-shadowing, the kind that we investigate in this study, is a type of retroactive interference where the introduction of IS after the CS interferes with the CS’s association with the US (sickness) (***Koen and Rugg, 2016***). Our results could be applicable to other forms of overshadowing and different paradigms of memory interference (***Polack et al., 2017***), as in all cases, a competition between reactivating assemblies would provide a neural mechanism for these behavioral findings of competition between stimuli for association with an outcome.

Prominent previous models that aim to account for overshadowing include the Rescorla-Wagner model and the Comparator hypothesis. The Rescorla-Wagner model explains overshadowing as a competition between stimuli for associative strength based on their salience and a prediction error during learning (***Rescorla and Wagner, 1972***). Another competing hypothesis is the Comparator hypothesis, where all associations are learned, but the expression of behavior depends on how the brain compares the associative strength of a target cue with other related cues at the time of retrieval (***Denniston et al., 2003***). Within the comparator hypothesis, overshadowing is not a competition of association, but due to interference at the time of retrieval (testing). Our work contributes to this theoretical landscape by offering a biologically plausible mechanism that could underlie the Rescorla-Wagner model–reactivation-based competition–that accounts for overshadowing through neural dynamics.

## Methods and Materials

### Network model

To study how credit is assigned in overshadowing, we simulate a recurrent network of adaptive exponential leaky integrate-and-fire neurons (***Brette and Gerstner, 2005***). We use this particular model with adaptation so that the network does not get stuck reactivating only one memory (generally the one with favorable initial conditions). Adaptation enables the network to reactivate both ensembles, and the time spent reactivating a stimulus representation would then serve as a read-out of that stimulus’s association with sickness. The network dynamics are given by the following equations:

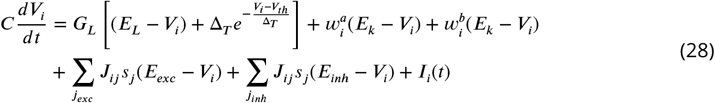

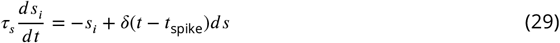

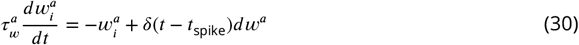

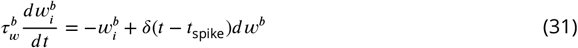

where *i* ∈ (1, *N* ). *V*_*i*_ is the voltage, *s*_*i*_ is the conductance and *w*_*i*_ is the adaption variable of the i th neuron. *V*_*th*_ is the threshold voltage and when the neuron exceeds the threshold, a spike is detected at *t* = *t*_spike_ and the voltage is reset to *V*_reset_. *ds* and *dw*^*a,b*^ are the increments in *s* and *w*^*a,b*^ with each spike.

*J* is the connectivity matrix. *J*_*ij*_ describes the synaptic weight between the presynaptic neuron *j* and the post-synaptic neuron *i*. The weights are selected such that there are two clusters of higher synaptic weights within the excitatory neurons. Within the cluster, the weights have the same value; however, the network behavior remains unchanged even when using a distribution of weights centered on the same mean. We assume there are no overlaps between the CS and the IS representations for most simulations (but see Supplementary Figure refoverlap). In contrast, taste representations are highly overlapping. Most neurons respond to all tastes at some point in time (***Fletcher et al. (2017***), unpublished data from Hannah Germaine, Katz lab). From a modeling perspective, we could focus only on neurons that are non-overlapping in both ensembles. This would be equivalent to what we have now.

We multiply the resultant connectivity matrix by an Erdős–Rényi model (***Erdös and Rényi, 1959***) to introduce sparseness and randomness. The Erdős–Rényi model constructs a matrix by connecting nodes (in this case, neurons) randomly. Each edge (synapse) is included in the connectivity matrix with probability *p*, independent of every other synapse. Table 2 lists the connectivity matrix parameters.

**Table 1.**
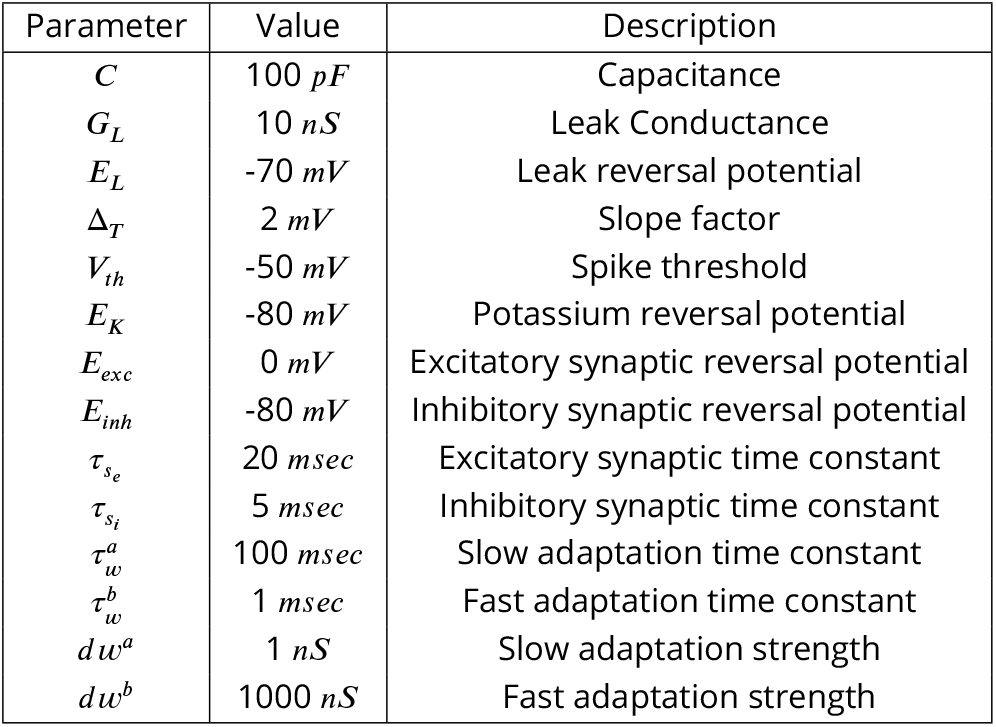
Parameter values used for simulations.

**Table 2.**
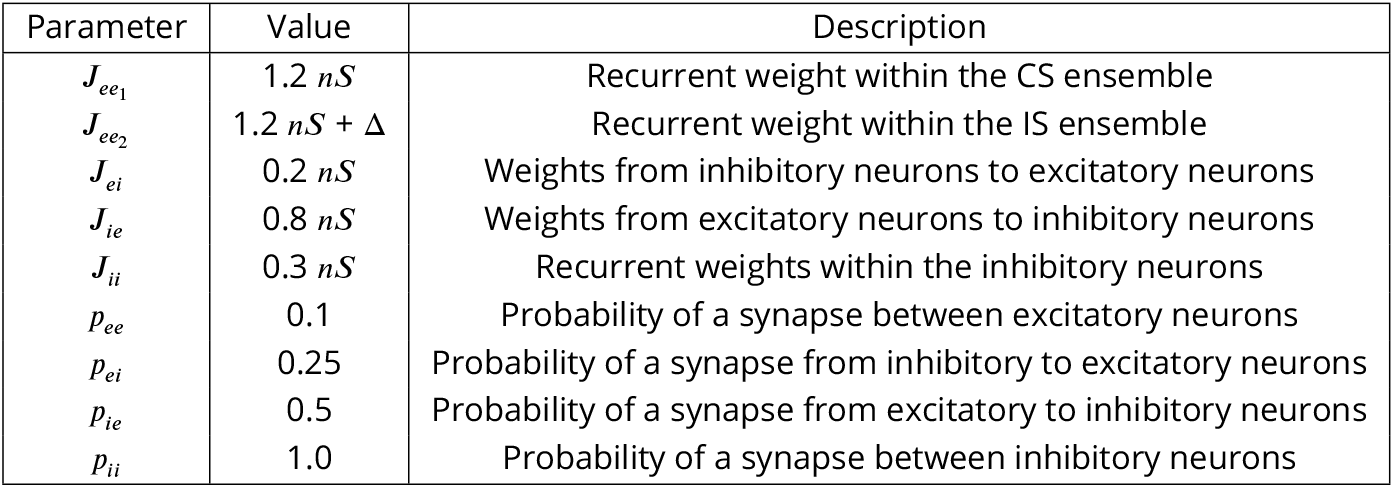
Parameters used for the connectivity matrix.

All neurons receive independent Poisson inputs, but for the excitatory neurons, the neurons share the rate of the Poisson processes, which is given by an OU process (See Figure methods C). The Poisson inputs are then converted into conductances, and Table 3 lists the parameters for the input current. We employ an OU process for rate modulation, allowing temporally extended upticks in the inputs that neurons receive, which enables reactivation events in the network.

**Table 3.** Parameters for the stochastic inputs.

| Parameter | Value | Description |
| --- | --- | --- |
| $\tau_{noise_e}$ | 2 msec | Time constant for excitatory noise |
| $\tau_{noise_i}$ | 5 msec | Time constant for inhibitory noise |
| $g_{noise_e}$ | 0.6 nS | Excitatory noise conductance |
| $g_{noise_i}$ | 0.5 nS | Inhibitory noise conductance |
| $r_{noise_{ee}}$ | 4500 Hz | Rate of noise for *ee* connections |
| $r_{noise_{ei}}$ | 1000 Hz | Rate of noise for *ei* connections |
| $r_{noise_{ie}}$ | 1000 Hz | Rate of noise for *ie* connections |
| $r_{noise_{ii}}$ | 1000 Hz | Rate of noise for *ii* connections |

We model short term synaptic plasticity: facilitation and depression. The dynamics of the facilitation variable (*F* ) is given by

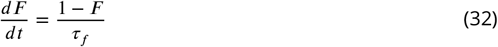

Following each spike the variable is updated via:

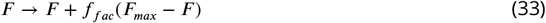

where *f*_*fac*_ denotes the degree of facilitation, *F*_*max*_ is the maximum value *F* can take and *τ*_*f*_ is the time constant for decay of *F* . Dynamics of the depression variable *D* is given by:

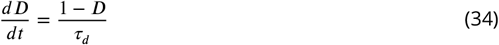

Following each spike the variable is updated via:

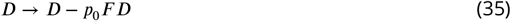

Where *p*_0_ is the initial release probability of the synaptic vesicles, *F* is the facilitation variable and *τ*_*d*_ is is the time constant for decay of *D*.

### Reactivation detection

When we simulate the network with correlated Poisson inputs, we observe reactivation of both tastes, similar to that observed in the gustatory cortex (Fig. react). We consider a stimulus to be reactivated if the ensemble average exceeds 10 Hz. We calculate the fraction of time the network spends reactivating each ensemble as a readout of its association with sickness: the more time the network spends reactivating a stimulus during sickness, the more it will be associated with sickness.

### Competition Index

As we propose, the amount of reactivation during malaise is monotonic in the degree of conditioning. Competition shapes the credit assignment for the two stimuli. Therefore, we aimed to quantify the degree of competition. ***Kuhl et al. (2011***) shows that competition between memories (stimulus representations) can be measured by the relative degree of reactivation. We calculate the fraction of time the network spends reactivating the CS and the IS as a function of Δ. To capture how the IS reactivation affects the CS reactivation, we look at the ratio of slopes of the lines representing the time spent in the CS and IS as a function of Δ.

Competition between the IS and the CS could occur due to a limited number of reactivations or a limited time for reactivation. In a limited-time competition, the time network spends reactivating one stimulus reduces the time available for the other stimulus to be reactivated. We refer to this as the competition due to the finite time available because it involves competing for time, and we want to remove the time factor in the cases where a lot of time is spent in a state, to investigate how different parameters affect the competition for reactivation between the two ensembles.

The degree of finite time competition is found as follows:

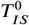 is the fraction of time spent by the network in the IS state at Δ = 0 and 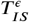 is the fraction of time spent by the network at a non-zero value of Δ = 𝜖. At Δ = 0, the network spend equal amount of time in the CS and the IS states.

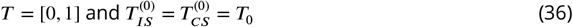

The network at Δ = 𝜖, spends additional time in the IS state, and as the maximum value of total time *T* is one, the fraction of time available for CS to be reactivated is

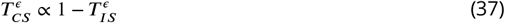

Similarly,

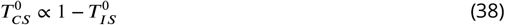

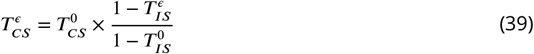

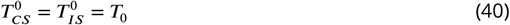

The degree of competition calculated as ratio of slopes due to time competition:

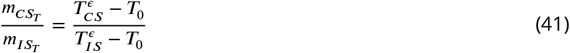

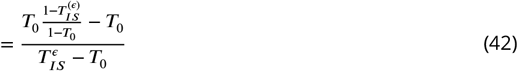

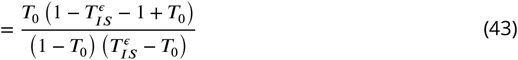

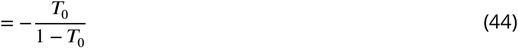

The competition index is then given by:

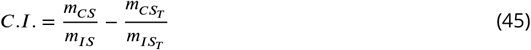

Where *m*_*CS*_ and *m*_*IS*_ are the slopes found by fitting lines to the time spend in state curves for CS and the IS.

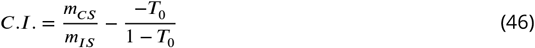

